# Application of long-read sequencing for genotyping, epigenetic profiling and surveillance of *Yersinia pestis* isolates from natural foci and disease outbreaks in Central Asia

**DOI:** 10.1101/2025.10.14.682323

**Authors:** Aigul A. Abdirassilova, Duman T. Yessimseit, Altyn K. Rysbekova, Altynai K. Kassenova, Beck Z. Abdeliyev, Zauresh B. Zhumadilova, Gulnara Zh. Tokmurziyeva, Arnat S. Dikhanbayev, Sanzhar D. Agzam, Vladimir L. Motin, Oleg N. Reva

## Abstract

This study explores the application of long-read sequencing technologies for genotyping, epigenetic profiling, and epidemiological monitoring of *Yersinia pestis* isolates obtained from natural foci in Central Asia and previous zoonotic outbreaks. Computational tools for genome assembly and genotyping were developed, enabling high-precision identification of both chromosomal and plasmid sequences, including the small cryptic pCKF plasmid. SNP-based genotyping distinguished the major *Y. pestis* biovars (Antiqua, Medievalis and non-main) and revealed cluster-specific diversity among Medievalis (MED) isolates, identifying a group of strains particularly prone to transmission from rodents to domestic animals and humans, which can be facilitated by the plasmid pCKF. Specific genomic polymorphisms were identified in sub-clades of MED isolates, which allow their identification with high precision. Additionally, comparative epigenomic analysis uncovered strain-specific cytosine methylation patterns at *cg*GATCG motifs, which may be linked to genome function regulation and adaptation to different hosts and environments. These findings demonstrate the effectiveness of long-read sequencing technologies in revealing both genetic and epigenetic features of bacterial pathogens, contributing to our understanding of the evolutionary mechanisms underlying the emergence and spread of this especially dangerous infection.

## 1. Introduction

Advancements in microbial genome sequencing technologies over recent decades have opened fundamentally new opportunities for researchers to study bacterial evolution and adaptation mechanisms to environmental changes, both at the genetic and epigenetic levels. The progress in recent years has been driven by rapid improvements in the quality and reliability of sequencing using SMRT PacBio, Oxford Nanopore (ONT) and Side Nanopore Sequencing (NSS) technologies, which generates long DNA reads and, many of them are able to record nucleotide call kinetics. The kinetics data allow for the identification of methylated nucleotides and the associated motifs at the binding sites of methyltransferases (MTases) on the DNA molecule [1].

Advances in whole-genome sequencing provide a new approach to understanding the emergence of infectious disease outbreaks [2]. The World Health Organization (WHO) warns of a high likelihood of the emergence of both novel diseases and re-emerging infectious diseases that have acquired new characteristics, such as increased frequency of cross-species transmission between animals and humans, antibiotic resistance, or altered pathogenicity pathways [3-5]. The increased likelihood of new disease emergence is associated with climate change, overpopulation, and urbanization, which in turn lead to increased rates of migration and social conflict.

Plague is an example of an infectious disease that has caused three deadly pandemics, each affecting vast regions and tolled millions of deaths. Notably, each pandemic was characterized by a shift in transmission strategy and infection dynamics from the exclusively bubonic form of plague typical of the Justinian Plague, primarily transmitted from infected rats to humans via fleas, to the predominantly pneumonic form observed during the Manchurian outbreak in China at the beginning of XX century, in which the infection was easily transmitted from person to person. The Black Death of the Middle Ages represented an intermediate form, with both bubonic and pneumonic manifestations present [6-8].

The three major plague pandemics are associated with three main biovars of *Yersinia pestis*: Antiqua (ANT), Medievalis (MED), and Orientalis (ORI). In the literature, the abbreviations 2.ANT, 2.MED0, and 2.MED1 are commonly used. The prefix “2” denotes their placement within the branch 2 of the *Y. pestis* phylogenetic tree, distinguishing these modern clades of the ANT and MED biovars from ancestral and non-main lineages [9-11].

Some researchers consider these main biovars to be subspecies that diverged from an ancestral, non-main, low-virulence variant of the pathogen known as “*Y. pestis microtus*,” which strains were isolated from voles and pikas in remote mountainous regions of Caucasus, Central Asia, and China [12-14]. Several mutations have been identified that appear to have significantly increased the likelihood of transmission from rodents to humans. These include a frameshift mutation in a gene of the Rcs signalling pathway, which led to a substantial upregulation of the *hms* gene and, consequently, uncontrolled formation of a bacterial biofilm causing blockage in the esophagus of infected fleas [15,16]. This blockage hinders flea feeding making them to leave their hosts in search of a new blood source, thereby infecting other animals and humans. Additional mutations in the genes *ureD, napA, araC, ilvN*, and *yeaW* were specific to different biovars and likely also influenced pathogen virulence and transmission routes [17-20]. However, the complete picture of the genetic and metabolic changes in pathogenic *Y. pestis* strains across the three major clades distinguishing them from each other and from the non-classical clade remains unclear.

Since natural plague foci span vast areas across all continents between 55° northern and 40° southern latitudes where pathogens are regularly isolated from rodent populations, the evolution of the pathogen and the emergence of a new *Y. pestis* variant cannot be ruled out. Such a variant could differ in its transmission route from animals to humans and between humans, as well as in its pathogenicity mechanisms. An example of such evolutionary development is the recent discovery of the pCKF plasmid in *Y. pestis* strains from the Central Caucasus natural plague focus [21]. Although the role of this plasmid in virulence is still unclear, its spread through the *Y. pestis* population beyond the Central Caucasus region may indicate that the plasmid confers certain selective advantages to its carriers.

The role of epigenetic changes in the evolution of pathogens also remains unclear. DNA methylation by MTases involved in restriction-modification (RM) systems, paired with the activity of cognate restriction enzymes that cleave unmethylated DNA at the same recognition sites, serves as a form of bacterial immunity against viruses and conjugative plasmids [22]. At the same time, the most widespread form of methylation among Enterobacteriaceae is the methylation of adenine residues on both DNA strands at GATC motifs, controlled by the orphan Dam MTase [23]. Since the solitary Dam MTase is not associated with a restriction enzyme, it cannot protect the cell from viral invasion. It was therefore reasonable to assume that the widespread occurrence of Dam-mediated methylation in bacteria, mainly in enterobacteria but also in many other microorganisms, including phylogenetically distant Gram-positive bacteria, can be explained by the involvement of this DNA modification in the regulation of gene expression and in other functions important for maintaining genome stability [24-27]. The results of Dam methyltransferase suppression via experimental mutagenesis have been inconsistent. In some cases, mutant strains exhibited significant phenotypic changes [28], while in others no changes were observed [29]. Altogether, this points to the complex nature of epigenetic DNA modifications in the regulation of vital processes in bacteria.

Doubts about the role of Dam methylation in genome regulation can also been raised due to the high efficiency of this MTase, which methylates adenine residues at nearly 100% of GATC motifs regardless of cultivation conditions. It is difficult to envision how Dam MTase could influence cellular processes in the absence of variation in methylation patterns.

Recent discoveries shed light on the possible mechanisms by which Dam methylation influences gene regulation. It was found that GATC represents only the central core of more complex motifs, in which both adenine and adjacent cytosine residues are methylated. For example, in *Escherichia coli, Klebsiella pneumoniae*, and *Streptococcus pneumoniae*, methylation of adenine and cytosine occurs within the motifs *c*RGKG*at*C as well as in super-palindromes such as *c*RGKG*at*CMCY*g* [27, 30]. Hereafter, methylated bases and the bases opposite methylated residues on the complementary DNA strand are depicted in the motifs using lowercase italic letters.

While adenine methylation in the central part of the motif occurs with nearly 100% efficiency, cytosine methylation was detected in only 30–70% of motifs, making it plausible that this methylation can contribute to gene regulation. Indeed, recent publications have suggested that it is methylated cytosine, rather than adenine, that has a stronger influence on gene expression, thereby establishing a stable, strain-specific gene expression pattern [31, 32].

It was previously established that *Y. pestis* also belongs to the group of microorganisms with active Dam methyltransferase, which methylates adenine residues in G*at*C motifs throughout the genome [33]. Cytosine methylation was not investigated in that study.

This study builds on our previous use of Illumina short-read sequencing to genotype *Y. pestis* strains, which distinguished desert and upland subpopulations of 2.MED1 isolates in Central Asia [34]. That work showed high conservation of core genomes among MED isolates from Central Asia, rendering them indistinguishable by approaches such as MLST and MLVA. We hypothesize that third-generation long-read platforms, such as PacBio SMRT (Revio) and GSeq 500, can further improve the sensitivity and specificity of pathogen genotyping. However, using long reads required a complete reconstruction of the computational pipelines for quality control, assembly, and genotyping, owing to incompatibilities between tools designed for short- and long-read data.

Beyond enabling more contiguous assemblies and more reliable variant detection, the PacBio SMRT Revio platform also calls methylated nucleotides, adding an epigenetic layer to comparative analyses. Here, we apply these long-read technologies to characterize patterns of genetic variation and DNA methylation in *Y. pestis* collected from natural rodent reservoirs in Central Asia and during zoonotic outbreaks in domestic camels with transmission to humans. Our goal is to develop genotyping approaches that track pathogen spread in nature and identify sources of human and animal infections using advanced sequencing methods.

The *Y. pestis* strains analysed in this study were isolated over a broad time span, the earliest date back more than 60 years. Such temporal spread can potentially complicate genotyping because random mutations may accumulate during culture maintenance. To address this problem, we conducted a systematic search polymorphic loci that remain stable within lineages yet are discriminative across sources, prioritizing genetic polymorphisms for long-term molecular surveillance.

## 2. Methods

### 2.1. Yersinia pestis strains used in this study

*Yersinia pestis* strains used in this study were selected from the National Collection of MicroOrganisms (NCMO) at the National Scientific Center of Especially Dangerous Infections named after Masgut Aikimbayev (NSCEDI, https://nscedi.kz/en/) in Almaty, Kazakhstan. Strains were selected to represent different natural plague foci of Central Asian deserts and mountains, including isolates from infected domestic animals and human patients (Table 1).

**Table 1.** *Yersinia pestis* strains used in this study.

| Autonomous plague foci | Strain number | NCBI BioSample | Isolation year | Source of isolation |
| --- | --- | --- | --- | --- |
| Moyunkum | 28_YP48_MKUM | SAMN50285116 | April 4, 1973 | Libyan jird (*Meriones libycus*) |
| Pre-Balkhash desert | 66_YP38_PB | SAMN47876468 | September 09, 2017 | Great gerbil (*R. opimus*) |
| Volga-Ural sandy | 16_YP27_VU | SAMN50285117 | June 26, 2007 | Dead midday jird (*Meriones meridianus*) |
| Volga-Ural steppe | 15_YP93_VU | SAMN50285118 | June 11, 1981 | Unidentified fleas on ground squirrels (*Spermophilus pygmaeus*) |
| North Pre-Aral desert | 8_YP55_NP | SAMN47876430 | October 06, 2020 | Great gerbil (*R. opimus*) |
| North Pre-Aral desert | 21_YP93_NP | SAMN50285119 | October 14, 2023 | Great gerbil (*R. opimus*) |
| Mangyshlak | 23_YP48_MANG | SAMN50285120 | August 1, 2003 | Infected patient |
| Mangyshlak | 23_YP47_MANG | SAMN50285121 | August 17, 2003 | Camel (*Camelus bactrianus*) meat |
| Mangyshlak | 23_YP45_MANG | SAMN50285122 | July 30, 2003 | Infected patient |
| Mangyshlak | 23_YP50_MANG | SAMN50285123 | August 17, 2003 | Camel (*C. bactrianus*) meat |
| Pre-Alakol low mountains | 48_YP14_PAK | SAMN47876476 | August 21, 2003 | Great gerbil (*R. opimus*) |
| Sarydzhas high mountains | 31_YP22_SZ | SAMN50285124 | June 26, 1971 | Grey marmot (*Marmot baibacina*) |
| Pre-Ustyurt | 19_YP08_PrU | SAMN50285125 | August 5, 1967 | Bone marrow of a fallen camel (*C. bactrianus*) |
| Ustyurt | 20_YP78_UST | SAMN50285126 | July 25, 1968 | Infected patient |
| Ustyurt | 20_YP79_UST | SAMN50285127 | August 6, 1968 | Camel (*C. bactrianus*) |
| North-Aral | 21_YP14_NP | SAMN50285128 | August 27, 2003 | Camel (*C. bactrianus*) meat |
| Sarydzhas high mountains | 34_YP28_SZ | SAMN47876493 | August 6, 1973 | Fleas (*Rh. li ventricosa*) |
| Sarydzhas high mountains | 38_YP25_SZ | SAMN47876494 | June 20, 1983 | Grey marmot (*M. baibacina*) |
| Sarydzhas high mountains | 31_YP37_SZ | SAMN50285129 | June 20, 1983 | Grey marmot (*M. baibacina*) |
| Sarydzhas high mountains | 31_YP74_SZ | SAMN50285130 | June 14, 1950 | Grey marmot (*M. baibacina*) |
| Talas high mountains | 37_YP35_TLH | SAMN47876497 | August 28, 1980 | Unidentified fleas |

### 2.1 Nutrient media used for *Y. pestis* cultivation and diagnostics

The strains were cultivated on Hottinger liquid and agarous media for 48 hours at 28°C. Their cultural and morphological properties were studied following standard laboratory diagnostic methods described previously [35]. Colony formation was monitored using light microscopy after 12, 24, and 48 hours of cultivation. Another diagnostic medium used in this study was Hottinger agar with 1% blood hemolysate.

### 2.3. Diagnostic bacteriological tests and PCR analysis

Screening of the phenotypic properties and characteristics of the studied *Y. pestis* strains was conducted using traditional microbiological methods, including assessment of growth on nutrient media, enzymatic and denitrifying activity, and sensitivity to specific bacteriophages.

For fermentation activity testing, strains were cultured in 5⍰ml Hiss medium (1% peptone water, 1% Andrade indicator, pH⍰7.2) with 1% arabinose, rhamnose, or glycerol. Fermentation was indicated by a color shift from yellow to pink after 48⍰h at 28⍰°C.

For denitrification testing, a loopful of 24–48⍰hours *Y. pestis* culture was inoculated into 1⍰ml Hottinger broth (pH⍰7.2) with 0.1% KNO_3_ and incubated at 28⍰°C for 72⍰h. Addition of 0.5⍰ml Griess reagent produced a crimson color in positive cases.

In this study, two diagnostic phages were used: a broad-spectrum *Y. pseudotuberculosis* phage (Saratov, Russia; series 20, 04.2022–04.2025) and the *Y. pestis*-specific Pokrovskaya phage (series 21, 12.2022–12.2025) [36]. Phage sensitivity was assessed by the streak-and-spot method: *Y. pestis* strains grown at 28⍰°C for 48⍰h were streaked on Hottinger agar, and phages were spotted nearby [37]. Plates were incubated at 28⍰°C for 15–18⍰hours. *Y. pestis* shows no growth at phage spots, while *Y. pseudotuberculosis* is lysed only by the broad-spectrum phage.

A set of primers shown in Table 2 was recommended for *Y. pestis* biovar differentiation [38]. PCR was performed in a 25⍰µl reaction containing 5⍰µl of 5X Genta PCR master mix (TaqF polymerase, dNTPs, Mg^2+^, buffer), 1⍰µl each of forward and reverse primers (20⍰pmol/µl), 5⍰µl of template DNA, and 8⍰µl of nuclease-free water. Amplification was run on a Rotor-Gene Q thermocycler (QIAGEN) with an initial denaturation at 95⍰°C for 5⍰min, followed by 35 cycles of 95⍰°C for 35⍰s, 60⍰°C for 35⍰s, and 72⍰°C for 35⍰s, with a final extension at 72⍰°C for 10⍰min.

**Table 2.**
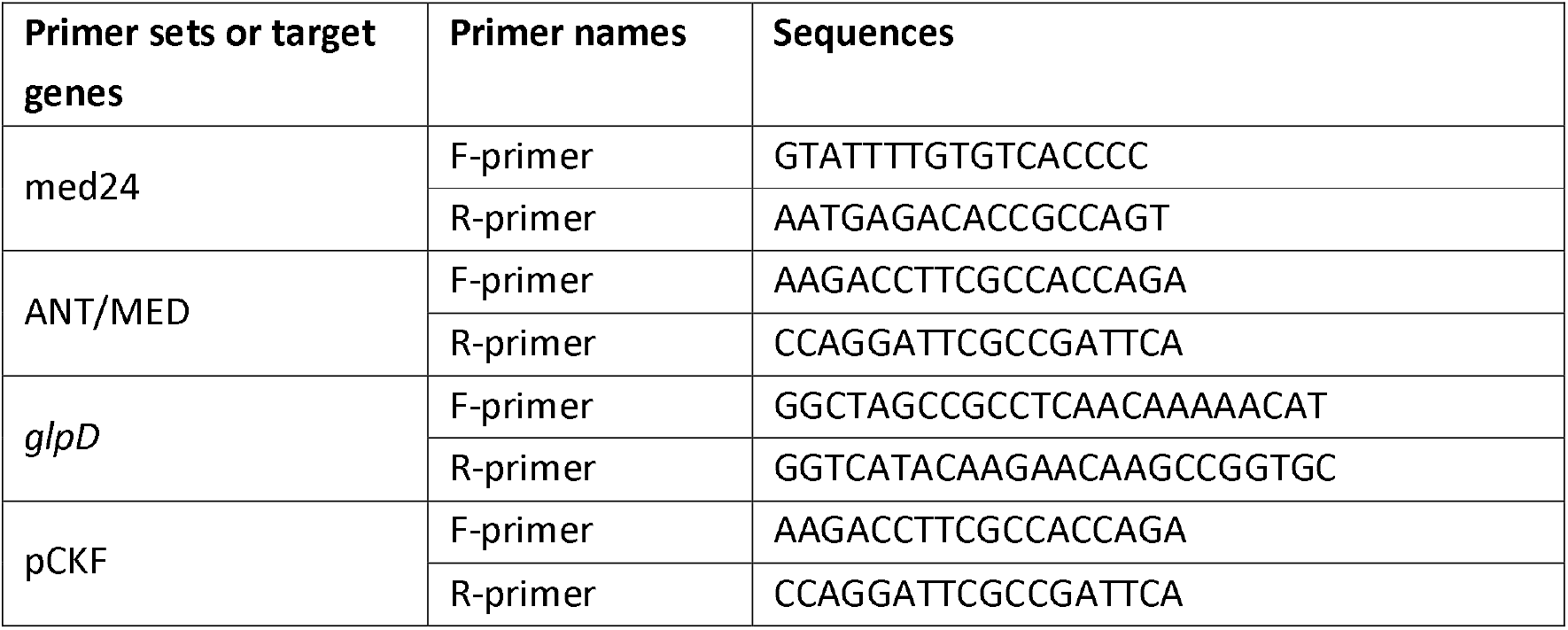
Diagnostic primers used in this study.

PCR products were separated on a 1.5% agarose gel stained with ethidium bromide and visualized under UV light using the UVP Mini Darkroom system. A Thermo Scientific GeneRuler 100⍰bp ladder was used for size estimation. Electrophoresis was performed on Amersham Pharmacia Biotech equipment.

### 2.4. DNA extraction and sequencing

Cell suspensions of *Y. pestis* strains were heat-treated at 100⍰°C for 10⍰min and centrifuged at 12,000⍰rpm for 2⍰min (Microlite RF centrifuge, Thermo). The supernatant was used for DNA extraction with the QIAamp DNA Mini Kit (QIAGEN, cat. no. 5⍰304), following the manufacturer’s protocol. DNA quality was evaluated by NanoDrop (260/280⍰nm ratio) and electrophoresis on a 1% agarose gel stained with ethidium bromide. All procedures were conducted under sterile conditions using disposable filter tips, gloves, and sterile tubes.

DNA sequencing was performed at the National Scientific Center of Especially Dangerous Infections on the long-read sequencers Geneus Gseq 500 (http://en.geneus-tech.com/proddetail.aspx?id=1) installed in the NCMO and two isolates representing 2.MED1 and 2ANT biovars were sequenced using the SMRT PacBio Revio platform provided by the Geneus Technologies (Almaty, Kazakhstan).

### 2.5 Computational pipeline LRASS for *Y. pestis* chromosomal and plasmid sequences

A computational pipeline was developed to assemble the *Y. pestis* chromosome and plasmids using SMRT PacBio long reads (Figure 1). This pipeline, implemented in Python 3.12.3, utilizes both *de novo* and reference-based assembly approaches, whose outputs are then combined into high-quality assembled sequences. The genome of *Y. pestis* SCPM-O-B-6899 (SAMN07176224), comprising one chromosome and four plasmids (pMT1, pCD, pPCP, and pCKF), sequenced using PacBio technology, was used as the reference sequence [21].

**Figure 1.**
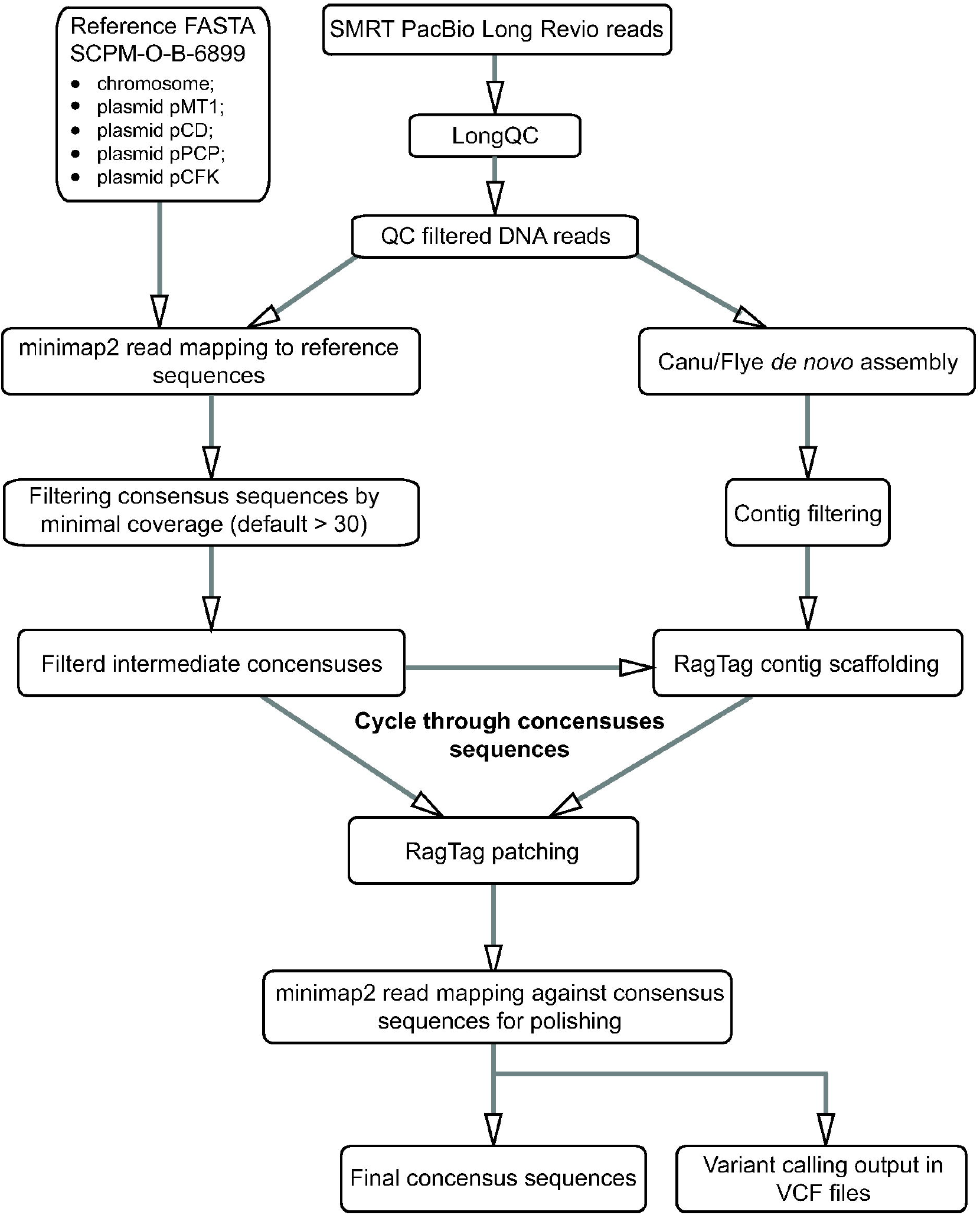
Y. pestis genome assembly pipeline LRASS (Long Read Assembler).

Processing of DNA reads begins with quality control and trimming using LongQC v1.2.0c [39]. The FASTQ files of trimmed reads and the FASTA file of reference sequences, in which replicons are represented as individual records, serve as input files for the LRASS (Long Read Assembler) pipeline, available for download from [https://zenodo.org/records/16571119]. For *de novo* assembly, Canu v2.0 [40] and Flye v2.9.6 [41] were used. DNA read mapping against the reference sequence was performed using Minimap2 v2.24-r1122 [42]. Consensus sequences were then extracted from the read alignments using Samtools v1.19.2 and BCFtools v1.19, both part of the HTSLIB 1.19 package.

*De novo* assembled contigs were scaffolded and patched using RagTag v2.1.0 [43], which employed replicon consensus sequences generated by aligning DNA reads to the reference sequences. The consensus replicon sequences produced by RagTag were filtered based on average sequence depth to avoid false assembly of plasmids not present in the current genome, using a coverage threshold of ≥30x. The replicon sequences were further polished through repeated alignment of the original DNA reads and extraction of consensus sequences and variant call files (VCFs) using BCFtools. The final whole-genome consensus sequences were annotated using Prokka v1.13.3 [44] and shuffled to ensure genome beginning from the replication origin near the *dnaA* gene on the direct strand.

An additional set of VCF files was generated by aligning the reads against the reference chromosome sequence of *Yersinia pseudotuberculosis* IP_2666pIB1 (CP032566) [45]. This strain was selected as a representative of the ancestral genome of *Y. pestis*, which was sequenced using similar PacBio technology. Single nucleotide polymorphisms (SNPs) predicted in the VCF files were filtered by quality and frequency using the following parameters: quality score (QUAL ≥ 30), depth of coverage (DP ≥ 10), mapping quality (MQ ≥ 40), mapping quality zero fraction (MQ0F ≤ 0.1), and the number of occurrences of the same polymorphism in the sequenced genomes (≥ 2).

For each identified SNP, an 81-bp context sequence centered on the polymorphic site was extracted from the reference and stored in a separate library file. Overlapping context sequences were merged. We developed an in-house Python script (https://zenodo.org/records/16572835) that uses BLASTN to query the assembled genomes with these context sequences, allowing up to six mismatches. The script outputs a binary (0/1) matrix for the analyzed *Yersinia* genomes, where 0 denotes a match of the context sequence to the reference sequence (putative ancestral state) and 1 denotes a mismatch (derived state). Polymorphic loci that were consistently observed in either state across minimum three genomes were retained as genotyping markers. This procedure improved the signal-to-noise ratio by filtering out spurious single-nucleotide variants (e.g., from sequencing errors or mutations arising during laboratory passaging). We then constructed a clustering dendrogram using the Camin-Sokal parsimony algorithm implemented in Bio.Phylo.TreeConstruction (Biopython v1.85).

Cytosine methylation in the assembled genomes was predicted using the program Jasmine v2.4.0 (https://github.com/PacificBiosciences/jasmine). Methylation motifs were identified using the MultiMotifMaker program (https://github.com/bioinfomaticsCSU/MultiMotifMaker) [46]. Statistical analysis and visualization of the distribution of modified cytosine residues were performed using the program SeqWord Motif Mapper v.3.2.6 (https://github.com/chrilef/BactEpiGenPro) [30]. This program performs statistical validation of the biased distribution of methylated or unmethylated motifs across coding regions, non-coding regions, transcriptional start codon (TSC) upstream regions, and genomic island inserts (GIs), which are automatically predicted by the program in bacterial genomes. The program also filters out sporadic, non-canonical methylation sites to retain nucleotides methylated within conserved canonical motifs. This filtering was important to reduce noise from unstable non-canonical sites that may arise in culture-collection strains due to differences in selective pressures between laboratory and natural environments.

### 2.6. Statistical validation of clade segregation by genetic polymorphisms

Clade segregation accuracy was assessed using the random forest classification algorithm, implemented via the *RandomForestClassifier* and *model_selection.cross_val_score* functions from the Python library *sklearn* (version 1.7.0). To evaluate whether the observed distribution of strains among clusters based on genetic polymorphism analysis significantly differs from a random distribution, a bootstrap analysis with 1,000 iterations was performed. In each iteration, the strains were randomly assigned to clusters, and classification accuracy scores were calculated. The likelihood that the accuracy from a random assignment equals or exceeds the actual score was estimated as a p-value using the *mean* function from the NumPy library (version 2.2.4).

The contribution of each polymorphic site to strain segregation was assessed by calculating p-values indicating the significance of cluster differentiation based on the given polymorphism, using the *chi2_contingency* function from the Python library SciPy (version 1.15.1). The Benjamini-Hochberg False Discovery Rate (FDR ≤ 0.05) correction for multiple testing [47] was applied to the calculated p-values using the function *multipletests* from the Python library *statsmodels* (version 0.14.4). This statistical validation of the separation power of individual polymorphisms and overall group separability was implemented in the in-house pipeline MatrixClassifier v1.1, available at https://zenodo.org/records/16618036.

Allelic states of the polymorphic sites were verified by whole genome alignment visualization using Mauve v.20150226 [48].

### 2.7. Availability of data and materials

All genome sequences generated in this study have been deposited in NCBI under BioProject PRJNA1249055 and the BioSample accession numbers provided in Table 1. The in-house pipeline scripts written in Python 3.12.3 are available from the Zenodo database at the following links: LRASS (Long Read Assembly pipeline) v1.0 (https://zenodo.org/records/16571119; DOI: 10.5281/zenodo.16571118); VARCALL (Yersinia Polymorphic Site Caller) v1.5 (https://zenodo.org/records/16572835; DOI: 10.5281/zenodo.16572835); MatrixClassifier v1.1 (https://zenodo.org/records/16618036; DOI: 10.5281/zenodo.16618036).

## 3. Results

In total, 21 *Yersinia pestis* strains were selected from the National Collection of Microorganisms (NCMO) at the National Scientific Center of Especially Dangerous Infections named after Masgut Aikimbayev (NSCEDI). These strains were chosen to represent different natural plague foci in Central Asia (Table 1), also taking into account the phenotypic diversity of the isolates discussed below.

### 3.1. Characterization of the selected *Y. pestis* isolates by phenotype and genotype

Laboratory test results for the selected *Y. pestis* isolates are presented in Table 3. All strains showed susceptibility to lysis by both *Y. pestis*-specific and *Yersinia* broad-range bacteriophages, confirming *Y. pestis*. One isolate 37_YP35_TLH obtained from the Talas high-mountain plague focus, was capable of fermenting rhamnose, unlike the other isolates. All isolates fermented glycerol, and most were also positive for arabinose fermentation, with the exception of the strain 37_YP35_TLH from the Talas mentioned above. Variation was also observed in nitrate reduction capacity, with five isolates testing nitrate-positive that is characteristic for the ANT biovar, while the rest were negative.

**Table 3.**
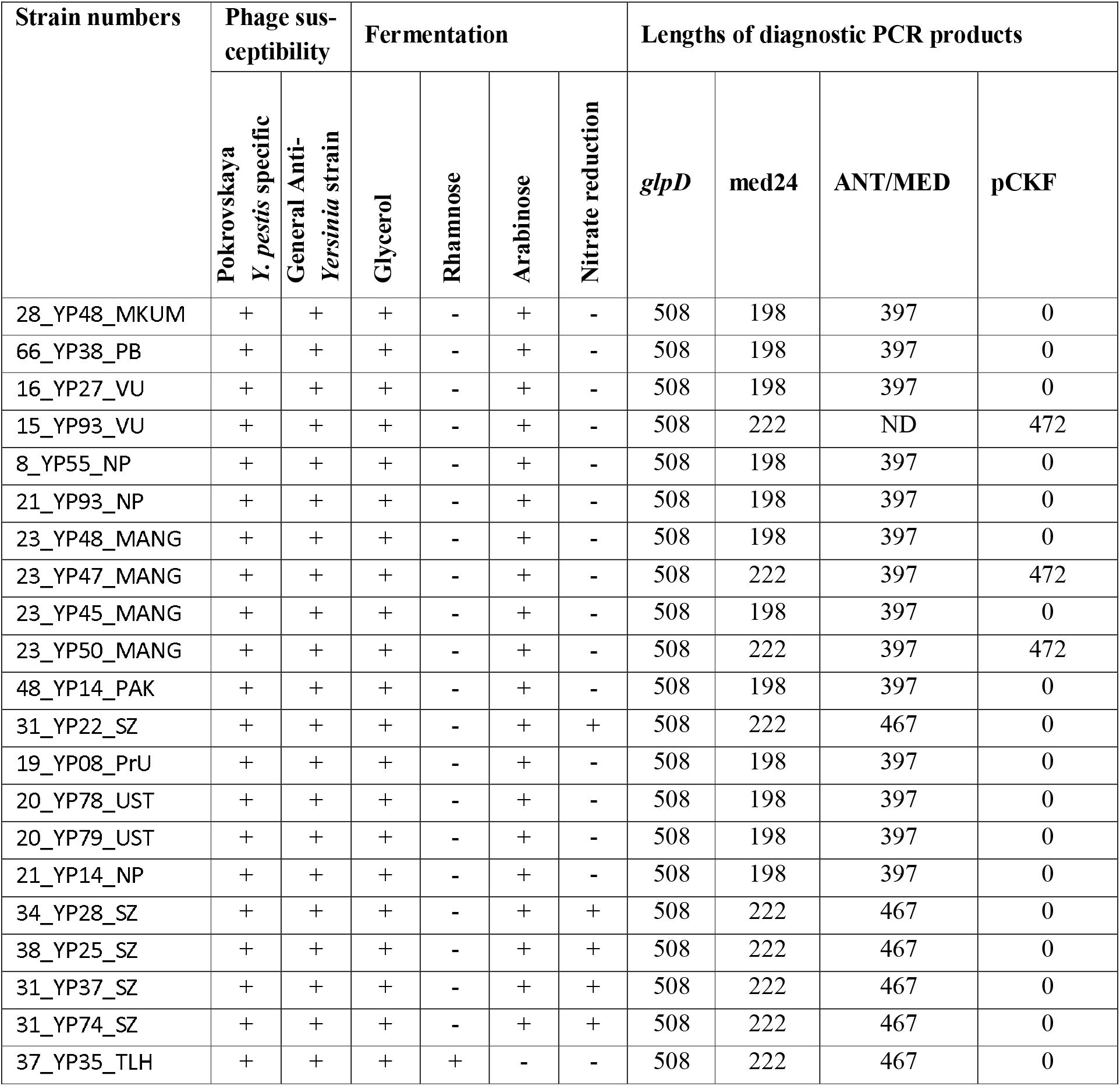
Biochemical tests and phage susceptibility.

PCR assays targeting polymorphic regions within the *rpoB* gene did not detect the deletion typically associated with the Orientalis biovar (data not shown). Likewise, analysis of the *glpD* gene did not reveal sequence variation among the studied isolates and confirmed the absence of the strains of the Orientalis biovar (Table 3). When tested with MED-specific primers (med24 and ANT/MED), most isolates amplified 397 bp fragments consistent with the MED clade. However, strains isolated from the high mountain regions of Sarydzhaz and Talas, including 31_YP22_SZ, 34_YP28_SZ, 38_YP25_SZ, 31_YP37_SZ, 31_YP74_SZ and 37_YP35_TLH yielded longer PCR products characteristic of the ANT and non-main *Y. pestis* biovars.

### 3.2. Identification of polymorphic sites suitable for distinguishing *Y. pestis* clades and biovars

Availability of newly sequenced genomes allowed identification of new polymorphic sites for distinguishing between *Y. pestis* lineage and sub-lineage. In total, 89 polymorphic nucleotide positions exhibiting different allelic states in at least two selected *Y. pestis* chromosomes were identified (Suppl. Table S1). The allelic states at these positions were converted into a 0/1 matrix, where 0 represents identity with the corresponding allele in the reference genome of *Y. pseudotuberculosis* IP2666pIB1 (CP032566), and 1 indicates a mutated allelic state. This matrix was used to construct a genome clustering dendrogram using the Camin-Sokal parsimony algorithm. In addition to *Y. pseudotuberculosis* IP2666pIB1, two other reference strains were included: *Y. pestis* SCPM-O-B-6899, isolated from a marmot in Kyrgyzstan in 1947 and maintained in Russian plague laboratories as a representative of the Antiqua (ANT) biovar; and *Y. pestis* SCPM-O-B-6530 isolated from *Citellophilus⍰tesquorum* fleas collected at the entrances to gopher (*Spermophilus musicus*) burrows in the Central-Caucasian high-mountain plague focus of Kabardino-Balkar Republic (Russia), in the year 2000 [21, 49]. Both genomes were sequenced using the SMRT PacBio technology. The latter strain, *Y. pestis* SCPM-O-B-6530, was identified as a representative of the MED0 lineage, which is considered as a basal clade within the Medievalis (MED) biovar [50, 51]. This strain also carries the rare cryptic plasmid pCKF.

The resulting dendrogram is shown in Figure 2. It is important to emphasize that the constructed dendrogram should not be interpreted as a phylogenetic tree, as it is not based on a computational model estimating evolutionary time. Moreover, it should not be excluded, that the higher level of similarity in allelic states may result from homoplastic adaptation to similar environmental conditions rather than from common ancestry [34, 52]. The purpose of this analysis was to develop an approach for grouping closely related organisms and distinguishing between groups to facilitate, in particular, the surveillance of different clonal lineages of the pathogen in nature.

**Figure 2.**
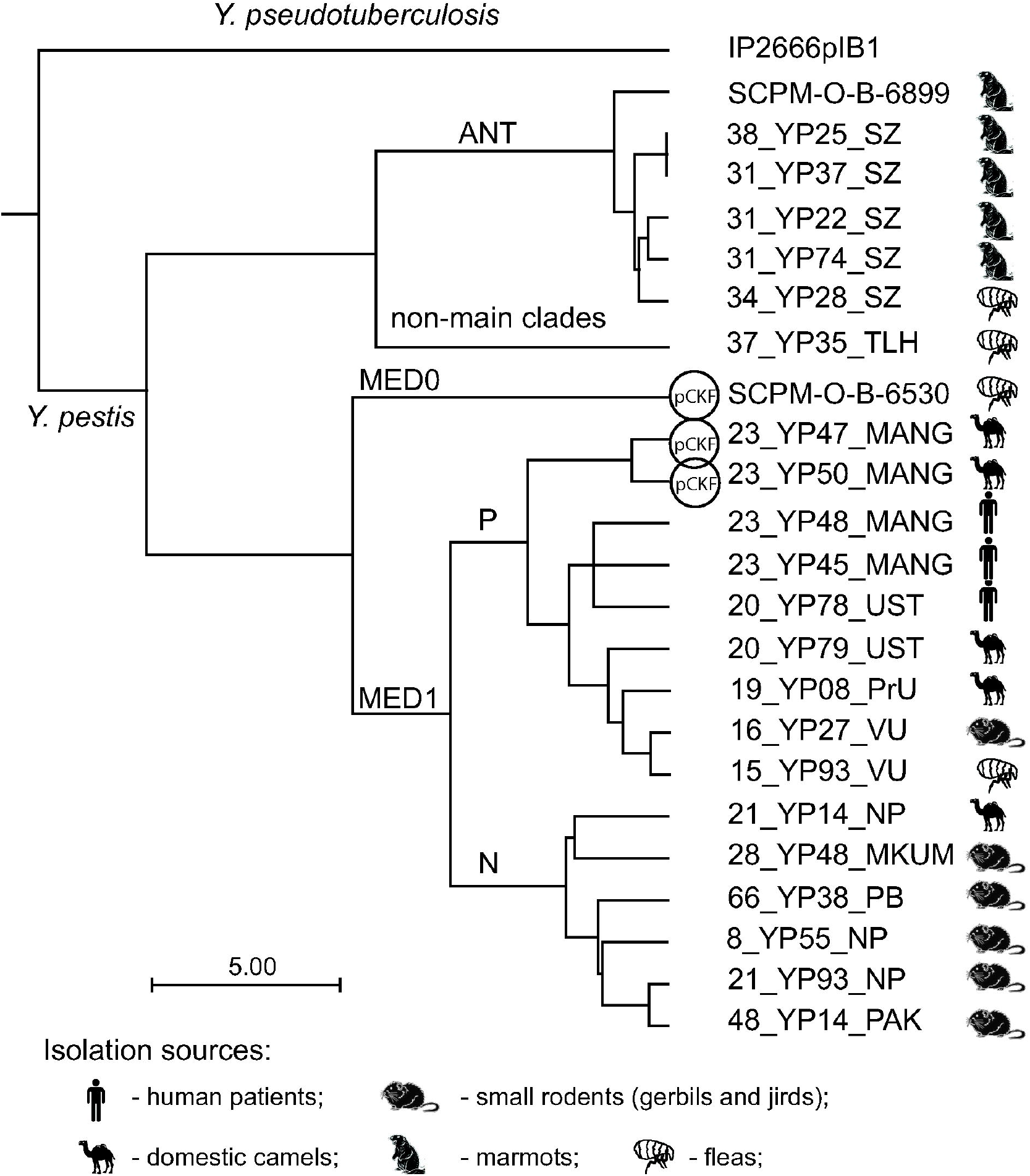
Dendrogram showing the similarity of allelic states at 89 polymorphic sites in the selected *Y. pestis* isolates and three reference genomes – *Y. pestis* SCPM-O-B-6530, *Y. pestis* SCPM-O-B-6899 and *Y. pseudotuberculosis* IP_2666plB1. Isolation sources and the strains caring the pCKF plasmid are indicated by respective icons. Branches of the tree leading to Antiqua (ANT), Medievalis (MED0 and MED1) and non-main clades, and additionally sub-clades P and N of the clade MED1, are labeled accordingly.

This comparison of polymorphic sites clearly separated MED1 *Y. pestis* isolates from isolates collected in the Sarydzhaz high-mountain plague focus, which belonged to the ANT biovar, along with a non-main clade isolate 37_YP35_TLH (Figure 2). The strain SCPM-O-B-6530 representing the MED0 biovar, is also clearly separated.

The MED1 isolates are further divided into two distinct sub-clades. Members of one sub-clade were isolated predominantly from small rodents inhabiting natural plague foci, whereas many members of the second sub-clade, which are centered near strain 15_YP93_VU, were associated with disease outbreaks among domestic camels, with subsequent transmission to humans. In Figure 2, these two sub-clades are labeled N (for natural plague foci) and P (prone to outbreaks), respectively.

Two strains, 23_YP47_MANG and 23_YP50_MANG, isolated from the meat of domestic camels that had developed plague infection, carried the small plasmid pCKF. This plasmid was previously reported in *Y. pestis* isolates from the Central Caucasus region, exemplified here by the reference strain SCPM-O-B-6899 [21]. Despite the sequence identity of the pCKF plasmids isolated in the Caucasus and Kazakhstan, the host strains differ significantly and belong to different biovars MED0 and MED1. This suggests that the plasmid can circulate within the *Y. pestis* population independently of the genetic background of the bacterial strains and within two distantly separated geographical regions.

Because the division of MED1 isolates into sub-clades with one of which appeared prone to causing disease outbreaks among domestic animals and humans is of epidemiological importance and has not been reported before, it is essential to validate this observation statistically and analyze the differences in detail. The significance of segregation between these two sub-clades of *Y. pestis* MED1 genomes was assessed using a random forest classifier, which achieved perfect classification accuracy of 1.0. Bootstrap analysis of clustering robustness confirmed the separation of strains between the two sub-clades based on the selected 89 polymorphisms, with a p-value of 0.0 (Suppl. Figure S1).

Analysis of the contribution of individual polymorphisms to this separation revealed four sites with p-values significantly lower than 0.01, which passed the Benjamini–Hochberg False Discovery Rate (FDR ≤ 0.05) correction. Three additional polymorphic sites showed statistically significant separation (p ≤ 0.05), but did not pass the FDR correction, for reasons explained in Table 4.

**Table 4.** Polymorphic sites with statistically reliable contribution towards separation between P and N sub-clades of *Y. pestis* MED1 isolates.

| Polymorphic site number* | Location in 15_YP93_VU | p-value | FDR p-value | Annotation |
| --- | --- | --- | --- | --- |
| 8 | 4,187,681 – 4,187,848 | 0.0009 | 0.019 | Insertion of IS100kyp transposone downstream of *osmY* in all strains of sub-clade N. |
| 24 | 3,191,545 - 3,191,575 | 0.0009 | 0.019 | Deletion of 37 bp at the beginning of *fabG_4* 3-oxacyl-reductase in all strains of sub-clade N. |
| 21 | 3,387,130 – 3,187,163 | 0.0009 | 0.019 | Deletion of 33 terminal bp of *aroG* phosphor-2-dehydro-3-deoxyheptonate aldolase in all strains of sub-clade N. Genome 20_YP78_UST of sub-clade P has similar truncation of this gene but due to an alternative mutation. |
| 48 | 1,622,050 - 1,622,100 | 0.0009 | 0.019 | Truncation of *yodF* symporter due to an insertion of ISEc39 transposone in all strains of sub-clade N. |
| 10 | 4,095,520 – 4,095,567 | 0.0151 | 0.269 | Deletion of 47 bp in the middle part of a hypothetical protein in all strains of sub-clade N, and also in the strains 23_YP45_MANG and 23_YP48_MANG of the sub-clade P. |
| 25 | 3,089,463 – 3,090,242 | 0.0239 | 0.303 | Disruption of MFS-type *ycaD_3* transporter by an insertion of an IS100kyp transposone in all strains of sub-clade P except for 16_YP27_VU. |
| 22 | 3,301,183 - 3,301,387 | 0.0236 | 0.303 | Highly polymorphic regions with several nucleotide deletions and substitutions between *sseA* 3-mercaptopyruvate sulfurtransferase and *ddc_2* L-2,4-diaminobutyrate decarboxylase. |

\* - polymorphic site numbers are given as in Suppl. Table S1.

### 3.3 Cytosine methylation in *Y. pestis* genomes

Recording the base call kinetics parameters in SMRT PacBio output sequence files enables the prediction of epigenetically modified bases. The program Jasmine used in this study is designed to predict methylated cytosine residues in PacBio HiFi reads generated by the SMRT PacBio Revio sequencer. The MultiMotifMaker program [46] predicted two major motifs, *c*GGATCG and CGATC*c*G, where cytosine methylation occurs in two sequenced strains.

Both of these motifs correspond to the same sequence, *cg*GATCG, which is associated with GATC sites that are stably methylated at adenine residues in *Y. pestis* genomes by the DAM MTase [33]. The distribution of methylated cytosine residues across the chromosomes of two *Y. pestis* strains, 31_YP22_SZ (ANT) and 15_YP93_VU (MED1), sequenced using Revio technology, was analyzed and statistically validated using the program SeqWord Motif Mapper [30]. The graphical outputs of the analysis are presented in Figure 3.

**Figure 3.**
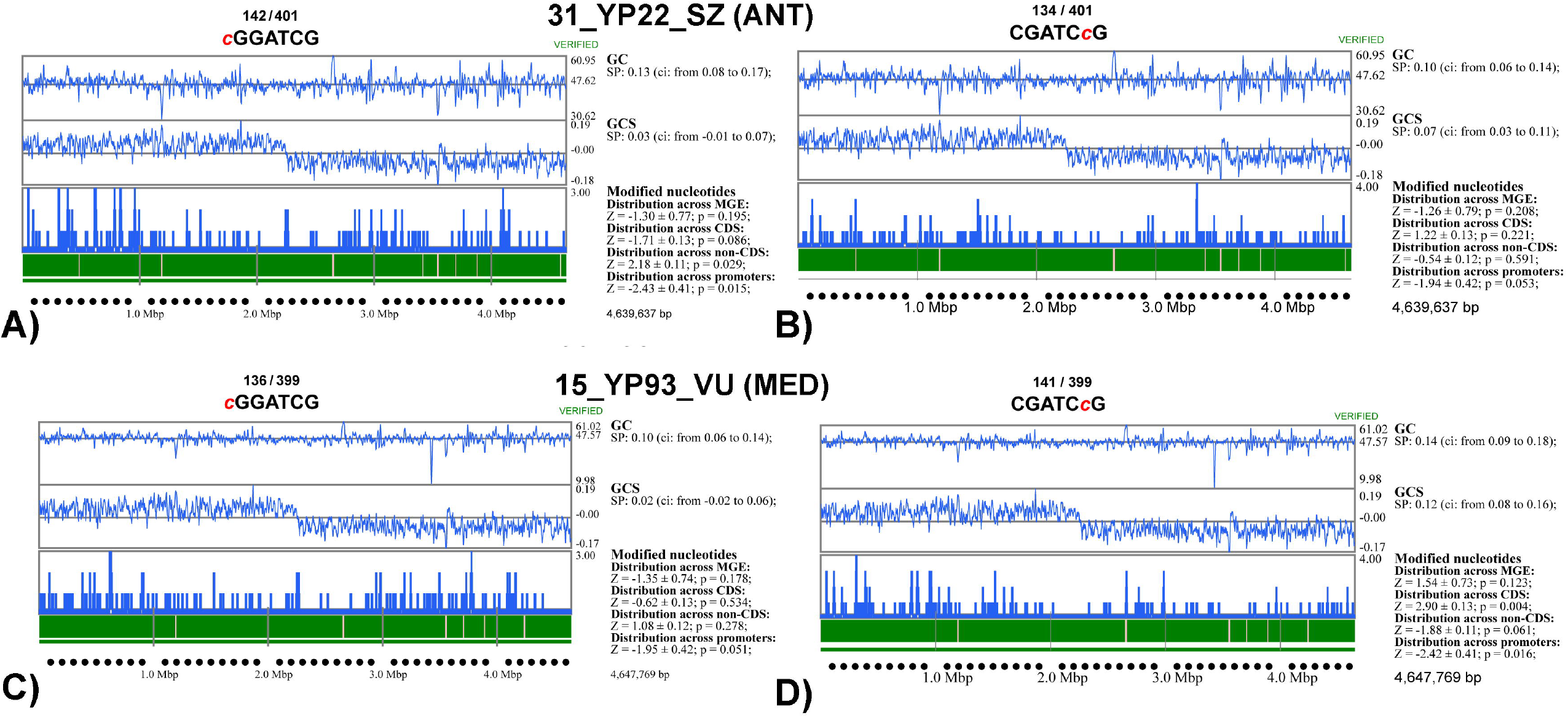
Distribution of methylated cytosine residues across *Y. pestis* chromosomes visualized using the SeqWord Motif Mapper program. A) methylation at cGGATCG motifs on the chromosome of *Y. pestis* 31_YP22_SZ representing the biovar ANT; B) methylation at CGATCcG motifs on the chromosome of *Y. pestis* 31_YP22_SZ; C) methylation at cGGATCG motifs on the chromosome of Y. pestis 15_YP93_VU; representing the biovar MED1; D) methylation at CGATCcG motifs on the chromosome of *Y. pestis* 15_VU93. The values *n*_*1*_ / *n*_*2*_ indicate the number of methylated motifs versus the total number of detected motifs in the sequence. The plots show GC content, GC skew, and methylation site density calculated using 8 kbp sliding windows with 2 kbp steps. Additionally, confidence intervals for Spearman correlations between GC content or GC skew and methylation frequencies within sliding windows, as well as Z-scores representing deviations in the frequency of methylated cytosine residues in coding regions, non-coding regions, 120 bp TSC-upstream regions, and horizontally acquired genomic regions, along with corresponding p-values indicating statistical significance.

The MultiMotifMaker program reported partial methylation of CGGATCG motifs, covering 35–37% of the available motifs in both genomes. The statistics on the biased distribution of methylated and unmethylated motifs across coding regions, non-coding regions, 120 bp TSC-upstream regions, and horizontally acquired GIs in the chromosomes of the two *Y. pestis* isolates are presented in Table 5.

**Table 5.** Biased distribution of methylated and unmethylated motifs across coding, non-coding 120 bp TSC-upstream and horizontally acquired regions in omes of two *Y. pestis* isolates sequenced using SMRT PacBio Revio technology.

|  | *Y. pestis* 31_YP22_SZ |  |  |  | *Y. pestis* 15_YP93_VU |  |  |  |
| --- | --- | --- | --- | --- | --- | --- | --- | --- |
|  | cGGATCG |  | CGATCcG |  | cGGATCG |  | CGATCcG |  |
|  | methylated | unmethylated | methylated | unmethylated | methylated | unmethylated | methylated | unmethylated |
| Coding | -1.70 ± 0.13<br>[p = 0.09] | -1.06 ± 0.1<br>[p = 0.29] | 1.28 ± 0.13<br>[p = 0.20] | 5.36 ± 0.09<br>[p = 0.0] | -0.43 ± 0.13<br>[p = 0.66] | -1.87 ± 0.1<br>[p = 0.06] | 2.97 ± 0.13<br>[p = 0.0] | 4.1 ± 0.1<br>[p = 0.0] |
| Non-coding | 2.16 ± 0.12<br>[p = 0.03] | 1.50 ± 0.08<br>[p = 0.14] | -0.59 ± 0.12<br>[p = 0.55] | -4.12 ± 0.08<br>[p = 0.0] | 0.92 ± 0.12<br>[p = 0.36] | 2.3 ± 0.08<br>[p = 0.02] | -1.94 ± 0.11<br>[p = 0.05] | -3.11 ± 0.08<br>[p = 0.0] |
| TSC-upstream | -2.40 ± 0.42<br>[p = 0.03] | -2.06 ± 0.3<br>[p = 0.04] | -1.92 ± 0.43<br>[p = 0.05] | -2.11 ± 0.28<br>[p = 0.04] | -1.94 ± 0.42<br>[p = 0.05] | -2.39 ± 0.3<br>[p = 0.02] | -2.41 ± 0.42<br>[p = 0.01] | -1.74 ± 0.31<br>[p = 0.08] |
| MGE | -1.81 ± 0.55<br>[p = 0.07] | 1.11 ± 0.57<br>[p = 0.27] | -1.77 ± 0.56<br>[p = 0.08] | 0.77 ± 0.53<br>[p = 0.44] | -1.90 ± 0.53<br>[p = 0.05] | 0.79 ± 0.53<br>[p = 0.43] | 2.20 ± 0.52<br>[p = 0.03] | -1.32 ± 0.54<br>[p = 0.19] |

It was found that the methylated motifs were well aligned with protein-coding sequences. CGATC*c*G motifs, both methylated and unmethylated, were more frequent in coding regions. The methylated cytosine residue was located downstream within the coding sequence relative to the methylated adenine in the G*a*TC motif. Conversely, methylation at *c*GGATCG motifs more frequently occurred on the reverse-complement non-coding DNA strand. A common feature for both types of motifs, whether methylated or not, was their significantly lower frequency in the 120 bp TSC-upstream regions. This finding suggests that mutations disrupting methylation motifs within regions potentially involved in transcription factor binding were beneficial and positively selected

Methylated sites were equally distributed between chromosomal replichores that is evidenced by absence of correlation with the GC-skew parameter. A weak positive correlation was observed between cytosine methylation frequency and GC content. This correlation can be explained by the more frequent occurrence of methylated sites within coding sequences, which typically have higher GC content compared to intergenic (linker) DNA. Methylated motifs were randomly distributed across the predicted horizontally acquired GIs.

## 4. Discussion

This study demonstrated the advantages of using SMRT PacBio long-read sequencing for genotyping, epigenetic profiling, and surveillance of *Y. pestis* isolates.

The genome assembly pipeline developed for this study [https://zenodo.org/records/16571119] enabled high-quality assembly of whole-genome sequences, including plasmid sequences, the number and content of which can vary between isolates. The designed library of genetic polymorphisms, along with the custom-developed analysis program [https://zenodo.org/records/16572835], demonstrated the suitability of this approach for genotyping, distinguishing between closely related strains, and monitoring the pathogens in natural environments. The ANT, MED, and non-main *Y. pestis* biovars were clearly separated in the dendrogram shown in Figure 2.

It should be emphasized that this dendrogram is not a phylogenetic tree. Rather, the approach was designed to differentiate between clades of *Y. pestis*, which are known to exhibit a high degree of genome conservation [11], for the purpose of pathogen surveillance and source identification in natural outbreaks.

The main focus of the developed *Y. pestis* genotyping program is on the MED biovar, which is dominant in Kazakhstan. Additionally, this study provided a clue to resolving the enigma of the zoonotic plague outbreaks reported in the Mangystau region in 2003. The strains 23_YP45_MANG and 23_YP48_MANG were isolated in clinics from patients, who were identified from domestic camels (*Camelus bactrianus*). The pathogens also were isolated from meat samples of the diseased camels (23_YP47_MANG and 23_YP50_MANG). Bacteriological testing confirmed that the isolates from the human patients and the camels belonged to the same MED biovar. However, further testing of the strains using PCR primers targeting the small cryptic plasmid pCKF revealed an unexpected difference: plasmid-specific bands were detected in the camel isolates but not in those from the human patients. Comparison of the patterns of genetic polymorphisms showed that the strains from the camels and the patients were similar but not identical. The plasmid-bearing isolates from the camels formed a tight cluster, distinct from other strains, whereas the human isolates clustered with other *Y. pestis* natural isolates from fallen camels, infected patients, and a dead midday jird (*Meriones meridianus*) from the same year, as well as a much earlier outbreak in 1968. All of these potentially pathogenic MED1 isolates were clearly separated into sub-clade (P in Figure 2) from natural *Y. pestis* MED1 strains (clade N) isolated from the great gerbil (*Rhombomys opimus*) and the Libyan jird (*Meriones libycus*), except for strain 3_YP14_ARAL, which was also isolated from camel meat in 2003.

Nevertheless, the strains belonging to these two sub-clades can be precisely distinguished by the polymorphic sites listed in Table 4. Several of these polymorphisms were associated with insertions of transposable elements, which are often characteristic of monophyletic groups of microorganisms [53]. The polymorphisms listed in Table 4 affect protein-coding genes, including: *fabG*_4 (a 3-oxoacyl-reductase involved in membrane fatty acid biosynthesis [54]), *aroG* (a phospho-2-dehydro-3-deoxyheptonate aldolase of the shikimate pathway), *yodF* (a putative symporter), *ycaD*_3 (an MFS-type transporter), and one large gene of unknown function.

The polymorphic region between the genes *sseA* and *ddc*_2 contains a microsatellite with multiple gttcatt and gctcatt repeats, which resemble hairpin-forming motifs typical of regulatory RNA secondary structures. However, this region has not been listed among predicted small RNA loci in *Y. pestis* genomes [55].

The plasmid pCKF is a small plasmid of approximately 5,400 bp, which, in addition to a replication-integration protein, encodes two type IV secretion system proteins and two hypothetical proteins [56]. The role of the plasmid remains unclear. In Kazakhstan, this plasmid was identified in strains that are genetically distant from the Central Caucasus isolates where the plasmid was initially identified (Figure 2), suggesting the possibility of unrestricted plasmid transmission within the *Y. pestis* population, regardless of lineage affiliation.

The isolation of different types of pathogens from domestic animals and infected humans suggests a stable mixed population infection, in which plasmid-bearing variants became dominant in camels without completely replacing plasmid-free variants of the pathogen, which became prevalent in humans. Similarly, plasmid pCKF was detected in isolate 15_YP93_VU by PCR using plasmid-specific primers; however, sequencing of this isolate did not reveal the presence of the plasmid. This discrepancy can be explained by the stringent assembly criteria in LRASS, which require a minimum local coverage of ≥30×. Such stringency can prevent assembly of plasmids not present in every sampled bacterial cell. We therefore conclude that the original culture was a mixture of plasmid bearing and plasmid free populations with the significant prevalence of the latter. The sequencing technique was not able to detect the plasmid in the subsequently isolated DNA while a more sensitive PCR method revealed the plasmid in the original sample obtained from the fleas. While the role of the plasmid pCKF in *Y. pestis* pathogenicity remains unclear, it is possible that the plasmid facilitates transmission of the pathogen from rodent hosts to domestic animals, which then serve as infection sources for humans. This could explain the increased number of plague outbreak incidents in the Caspian region of Kazakhstan in 2007, shortly after the first detection of the plasmid in the Central Caucasus plague focus in 2003.

An additional advantage of SMRT long-read PacBio sequencing is its ability to detect cytosine methylated nucleotides in the bacterial genome in parallel with genome sequencing. This study contributes to our understanding of the possible role of genomic DNA methylation in gene regulation and highlights the complexity of genome methylation mediated by DAM MTase. DAM is one of the most widely distributed MTases among enterobacteria and even some Gram-positive microorganisms, despite not being involved in RM systems that protect bacteria from phages.

Recent studies have suggested that this enzyme may play a role in gene regulation by controlling not only adenine methylation at G*at*C palindromes but also methylation of nearby cytosine residues. While adenine methylation occurs at nearly 100% of available sites regardless of growth conditions, cytosine methylation is more variable and condition-dependent. Methylated cytosine residues were significantly overrepresented in protein-coding sequences but absent from promoter regions. This pattern of cytosine methylation associated with DAM MTase has also been observed in other DAM-methylated bacteria such as *Escherichia coli, Klebsiella pneumoniae*, and *Streptococcus pneumoniae* [30]. In these organisms, cytosine methylation occurs within semi-conserved *c*RGKGatC motifs or even in super-palindromes like *c*RGKG*at*CMCY*g*. In *Y. pestis*, the methylation motif is more conserved, *cg*GATCG, and does not form super-palindromic structures, but otherwise aligns with the GATC-associated methylation of nearby cytosines reported across a wide range of bacterial species. Because DAM is often the only MTase found in bacterial genomes, it is likely responsible for both adenine and cytosine methylation in these motifs.

In both sequenced strains, 31_YP22_SZ and 15_YP93_VU, canonical cytosine methylation was strictly associated with the conserved motif *cg*GATCG. This conservation of canonical methylation and the absence of mutations within the *dam* genes in these two genomes demonstrates the functional stability of the associated DAM MTases in these strains, despite their prolonged cultivation under laboratory conditions (Table 1).

The comparison of cytosine methylation patterns in the genomes of two strains, 31_YP22_SZ (Suppl. Table S2) and 15_YP93_VU (Suppl. Table S3), representing the ANT and MED biovars respectively, revealed a striking difference, with only 50% overlap between methylated motifs located within protein-coding genes. For example, the gene *selB*, which encodes the selenocysteine-specific elongation factor, is methylated at three sites in 31_YP22_SZ, whereas none of these sites are methylated in 15_YP93_VU. Conversely, in 15_YP93_VU, the gene *ubiE* encoding ubiquinone/menaquinone biosynthesis C-methyltransferase is methylated at four sites, and the gene *rpoA* encoding subunit A of DNA-directed RNA polymerase is methylated at two sites; neither of these genes is methylated in 31_YP22_SZ.

This finding demonstrates significant diversification in patterns of cytosine methylation between these two strains. Further studies involving a larger number of *Y. pestis* strains are necessary to determine whether this diversification is driven by adaptation to different hosts and environmental conditions (ANT infections being associated with marmots in high-mountain regions, and MED infections being natural to gerbil and jird populations inhabiting the deserts of Central Asia), or by other factors.

## 5. Conclusions

This study demonstrated the value of long-read Geneus Gseq 500 and PacBio Revio sequencing for genotyping *Y. pestis* isolates, accurately tracing sources of infection, and revealing complex epigenetic patterns in bacterial genomes. Compared with recently published short-read approach [34], long-read platforms provided higher resolution for surveillance of closely related isolates. The “desert” MED genotype defined previously split into two groups, one of which appears particularly prone to transmission from rodent hosts to domestic camels and subsequently to humans. Our marker-selection strategy and publicly available tools for genome assembly and genotyping correctly grouped the *Y. pestis* collection, including strains maintained under laboratory conditions for more than 60 years. The genotyping system will be used to study introgression dynamics among *Y. pestis* subpopulations in Central Asia; an example analysis of introgressions between the “desert” and “upland” MED subpopulations was presented in our prior work [34].

Detailed analysis of the isolates from this group suggested a mixed infection involving two genetically similar but distinguishable variants of the pathogen, one of which carries the pCKF plasmid. The introduction of this plasmid from the Central Caucasus region into the Caspian regions of Kazakhstan in the 1970s may explain the recorded outbreak among farmers. The role of this small plasmid (∼5,400 bp) in pathogenicity and its distribution requires further investigation. Due to its small size, pCKF can be easily overlooked during genome assembly. However, the assembly pipeline proposed in this study, combined with PCR testing using plasmid-specific primers, enables effective detection and assembly of the plasmid in both pure and mixed cultures.

The novel sequencing approaches and computational tools provided insight into the complex nature of DAM-controlled genome methylation at adenine and cytosine residues. While adenine methylation is comprehensive and invariant, cytosine methylation adjacent to GATC motifs is strain-specific. This study demonstrated a statistically significant biased distribution of *cg*GATCG motifs and methylated cytosine residues across coding, non-coding, and TSC-upstream regions, as well as methylation pattern specificity in two *Y. pestis* isolates representing the ANT and MED biovars. However, the biological or evolutionary role of this methylation in *Y. pestis* remains unclear. It was hypothesized in the previous study, that the localization of methylated cytosine sites within motifs that may potentially form loops and junctions in mRNA molecules, their abundance in protein-coding regions, and their absence in putative promoter regions suggest that this type of methylation may influence mRNA turnover rather than transcriptional regulation [Lefebvre et al., 2025]. This hypothesis requires further investigation, involving transcriptomics, proteomics, and direct RNA sequencing using Oxford Nanopore technologies for studying RNA epigenetics.

## Acknowledgements

Authors acknowledge the company Microread Technology Ltd. (Almaty, Kazakhstan) and personally Mr. Yernazar Rashatbeck for help with installation of the sequencer at NCMO and provided training.

## Authors’ contributions

AAA contributed to project design, funding acquisition, bacterial strain collection, supervision, and manuscript drafting and finalization; YDT performed DNA extraction, sequencing, and manuscript drafting; RAK conducted molecular genetic studies; KAK was responsible for epidemiological studies and manuscript drafting; ABZ performed DNA extraction, sequencing, and manuscript drafting; ZZB contributed to conceptualization and supervision; TGZh was involved in conceptualization and manuscript editing; DAS managed culture collection; ASD carried out microbiological studies; MVI provided consultation and participated in manuscript drafting and finalization; RON performed genome assembly, bioinformatics, and programming, and contributed to manuscript drafting and finalization.

## Ethics approval and consent to participate

All work involving *Yersinia pestis* was conducted in full accordance with national biosafety regulations and international standards for handling BSL-3 agents. Routine surveillance, strain maintenance, and research activities are carried out under institutional biosafety protocols approved by the Biosafety and Ethics Committee of the National Scientific Center of Especially Dangerous Infections named after Masgut Aikimbayev (NSCEDI, https://nscedi.kz/en/). The use of *Y. pestis* strains for this study did not involve work with live animals or human subjects and was performed exclusively on archived bacterial cultures in compliance with national and institutional regulations.

## Supplementary materials

**Supplementary Figure S1.**
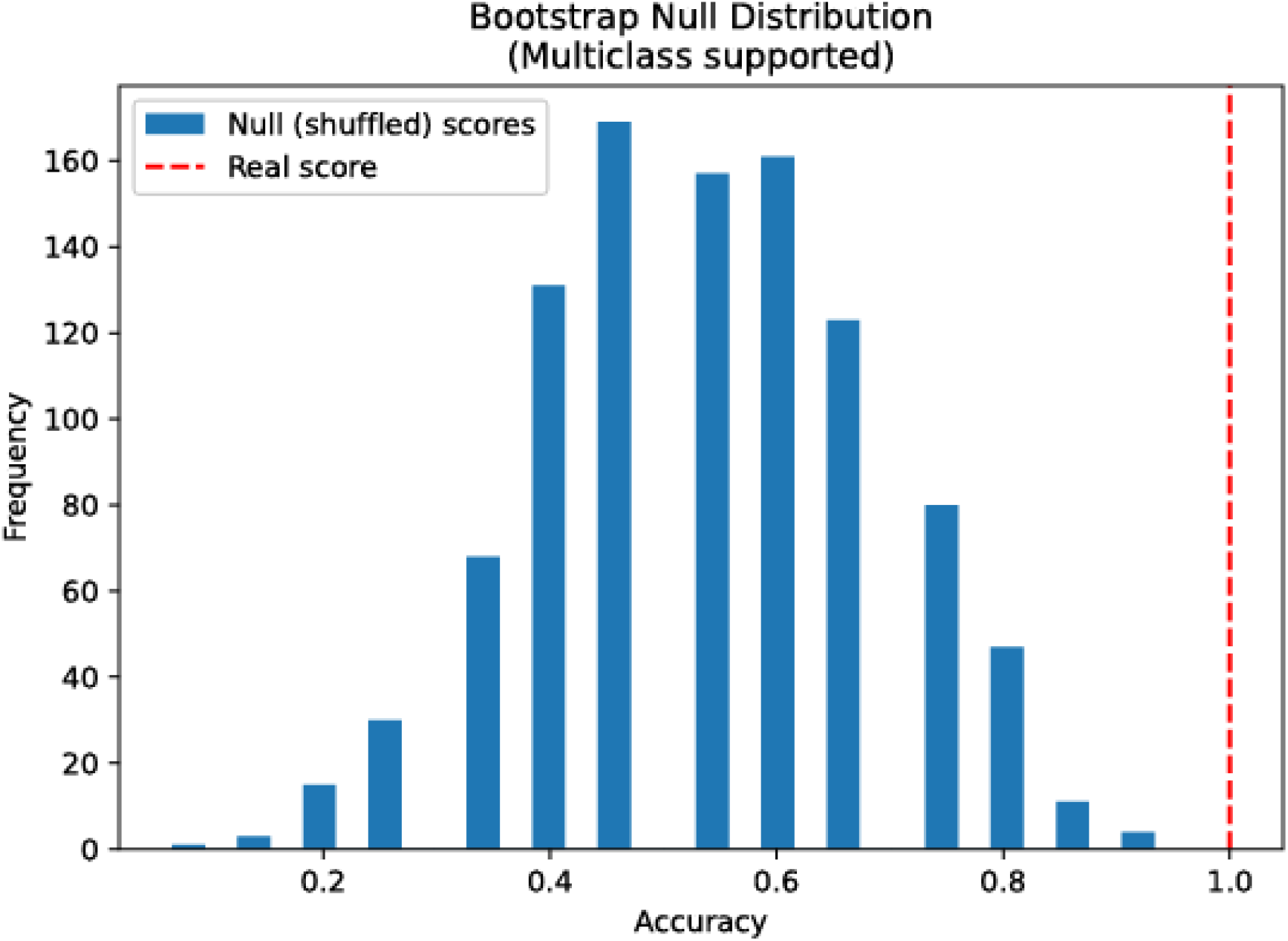
Bootstrap null distribution of classification accuracy across ‘N’ and ‘P’ groups of *Yersinia pestis* isolates. Histogram shows the distribution of classification accuracies obtained from 1,000 RandomForest runs with shuffled group labels (null distribution). The red dashed vertical line indicates the actual classification accuracy achieved using true group labels. The area of the histogram to the right of the real score represents the empirical p-value, estimating the probability of observing such classification performance by chance. This analysis supports the statistical significance of group separability based on the polymorphic features in the input matrix. Multiclass classification was supported and applied.

Supplementary Table S1. Matrix of polymorphic genetic loci. https://zenodo.org/records/17350388

Supplementary Table S2. Locations of methylated cytosine residues within cgATCCG motifs in the genome of Y. pestis 31_YP37_SZ. https://zenodo.org/records/17350388

Supplementary Table S3. Locations of methylated cytosine residues within cgATCCG motifs in the genome of Y. pestis 15_YP93_VU. https://zenodo.org/records/17350388

